# Ecological Interactions and the Netflix Problem

**DOI:** 10.1101/089771

**Authors:** Philippe Desjardins-Proulx, Idaline Laigle, Timothée Poisot, Dominique Gravel

## Abstract

Species interactions are a key component of ecosystems but we generally have an incomplete picture of who-eats-who in a given community. Different techniques have been devised to predict species interactions using theoretical models or abundances. Here, we explore the *K* nearest neighbour approach, with a special emphasis on recommendation, along with other machine learning techniques. Recommenders are algorithms developed for companies like Netflix to predict if a customer would like a product given the preferences of similar customers. These machine learning techniques are well-suited to study binary ecological interactions since they focus on positive-only data. We also explore how the *K* nearest neighbour approach can be used with both positive and negative information, in which case the goal of the algorithm is to fill missing entries from a matrix (imputation). By removing a prey from a predator, we find that recommenders can guess the missing prey around 50% of the times on the first try, with up to 881 possibilities. Traits do not improve significantly the results for the *K* nearest neighbour, although a simple test with a supervised learning approach (random forests) show we can predict interactions with high accuracy using only three traits per species. This result shows that binary interactions can be predicted without regard to the ecological community given only three variables: body mass and two variables for the species’ phylogeny. These techniques are complementary, as recommenders can predict interactions in the absence of traits, using only information about other species’ interactions, while supervised learning algorithms such as random forests base their predictions on traits only but do not exploit other species’ interactions. Further work should focus on developing custom similarity measures specialized to ecology to improve the *K*NN algorithms and using richer data to capture indirect relationships between species.

## 1 Introduction

Species form complex networks of interactions and understanding these interactions is a major goal of ecology [26]. The problem of predicting whether two species will interact has been approached from various perspectives [3, 22]. Williams and Martinez [32] for instance built a simple theoretical model capable of generating binary food webs sharing important features with real food webs [15], while others have worked to predict interactions from species abundance data [1, 7]. Being able to predict with high enough accuracy whether two species will interact given simply two sets of attributes, or the preferences of similar species, would be of value to conservation and invasion biology, allowing us to build food webs with partial information about interactions and help us understand cascading effects caused by perturbations. However, the problem is made difficult by the small number of interactions relative to non-interactions and relationships that involve more than two species [14].

In 2006, Netflix offered a prize to anyone who would improve their recommender system by more than 10%. It took three years before a team could claim the prize, and the efforts greatly helped advancing machine learning methods for recommenders [24]. Recommender systems try to predict the rating a user would give to an item, recommending them items they would like based on what similar users like [2]. Ecological interactions can also be described this way: we want to know how much a species would “like” a prey. Interactions are treated as binary variables, two species interact or they do not, but the same methods could be applied to interaction matrices with preferences. There are two different ways to see the problem of species interactions. In the positive-only case, a species has a set of preys, and we want to predict what other preys they might be interested in. This approach has the benefit of relying only on our most reliable information: positive (preferably observed) interactions. The other approach is to see binary interactions as a matrix filled with interactions (1s) and non-interactions (0s). Here, we want to predict the value of a specific missing entry (is species *x_i_* consuming species *x_j_*?).

Statistical machine learning algorithms [24] have proven to be reliable to build effective predictive models for complex data (the “unreasonable effectiveness of data” [17]). We will use a simple technique called the *K* nearest neighbour (*K*NN) algorithm both for recommendation (finding good preys to a species with positive-only information) and matrix imputation (filling a specific entry in a matrix with positive and negative interactions). The technique is simple: for a given species, we find the *K* most similar species according to some distance measure, and use these *K* species to base a prediction. For this study, we use a data-set from Digel et al. [11], which contains 909 species, of which 881 are involved in predator-prey relationships and 871 have at least one prey. The data comes from soil food webs and includes invertebrates, plants, bacteria, and fungi. In total, the data-set has 34 193 interactions. The data was complemented with information on 25 binary attributes (traits) for each species, plus their body mass and information on their phylogeny. We also briefly discuss a supervised learning method, random forests, which is used to predict interactions with only the species’ traits.

A summary of the three methods used can be found in table 1. The approaches are not directly comparable. For example, the positive-only *K*NN recommends preys to a species. If we remove a prey from a species, ask the algorithm to recommend a prey, and check whether the prey will come up as the recommendation, there are up to 881 possibilities. On the otherhand, the *K*NN algorithm with positive and negative values (matrix imputation) has to decide whether an entry is an interaction or a non-interaction, a 50% chance of success by random. These approaches have different uses. Positive-only algorithms are interesting because we are rarely certain that two species do not interact. Also, the *K*NN approach uses information on what similar species do, while random forests only rely on traits.

**Table 1:** Summary of the three methods used. The first two use the *K* nearest neighbour algorithm, with the Tanimoto distance measure in the positive-only case (recommendation), and the Euclidean distance with positive and negative values (matrix imputation). The Tanimoto *K*NN makes a recommendation, while the Euclidean *K*NN and random forests predict either an interaction or a non-interaction. The context for the Euclidean *K*NN and random forests are different: the former fills missing entries from a binary interaction matrix while the latter makes a prediction based only on the traits of the predator and its prey.

| Method | Input | Prediction |
| --- | --- | --- |
| Recommender (Tanimoto $KNN$) | Set of traits & preys for each species | Recommend new preys |
| Imputation (Euclidean $KNN$) | Interaction matrix with missing entries | Fill missing entries (0s and 1s) |
| Supervised learning (Random forests) | Traits (binary and real-valued) | Interaction (1) or non-interaction |

We show the *K*NN is particularly effective at retrieving missing interactions in the positive-only case, succeeding 50% of the times at recommending the right species among 881 possibilities. However, the *K*NN algorithm is significantly less accurate than random forests to predict an interaction with positive and negative data. With few traits, the random forests can achieve high accuracy (≈ 98% for both interactions and non-interactions) without any information about other species in the community. Random forests require only three traits to be effective: body mass and two traits based on the species’ phylogeny. Our results show that, with either three traits per species or partial knowledge of the interactions, it is possible to reconstruct a food web accurately.

## 2 Method

### 2.1 Data

The first data-set was obtained from the study of Digel et al. [11], who documented the presence and absence of interactions among 882 species from 48 forest soil food webs, details of which are provided in the original publication. 34 193 unique interactions were observed across the 48 food webs, and a total of 215 418 absence of interactions. For matrix imputation, we assume all entries in the 881 by 881 matrix which are not observed interactions are non-interactions. In order to improve representation of interactions involving low trophic levels species that were not identified at the species level in the first data-set, we compiled a second data-set from a review of the literature. We selected all articles involving interactions of terrestrial invertebrate species for a total of 126 studies, across these, a total of 1 439 interactions were recorded between 648 species. Only 88 absences of interactions were found. We selected traits based on to their potential role in consumption interactions (table 2). For each species or taxa, these traits were documented based on a literature review or from visual assessment of pictures. In addition to these traits, we included two proxies for hard-to-measure traits: feeding guild and taxonomy.

**Table 2:** The traits used. All traits are binary except for body mass, *Ph_0_,* and *Ph_1_.* We use taxonomy as a proxy of latent traits following [23]. To do so, we used the R package *ape* to obtain taxonomic distances between the species, perform classical multidimensional scaling (or principal coordinates analysis) [9] on taxonomic distances, and use the scores of each species on the first two axes (*Ph_0_* and *Ph_i_*) as taxonomy-based traits. These three real-valued variables are scaled to be in the [0,1) range. For the Tanimoto similarity index, these three continuous variables have to be converted to binary features. For each, we create four binary features (*n* = 881/4).

| Features | Abbr. | Description | $n$ |
| --- | --- | --- | --- |
| AboveGroud | *AG* | Whether the species live above the ground. | 538 |
| Annelida | *An* | For species of the annelida phylum. | 34 |
| Arthropoda | *Ar* | For species of the arthropoda phylum. | 813 |
| Bacteria | *Bc* | For species of the bacteria domain. | 1 |
| BelowGround | *BG* | For species living below the ground. | 464 |
| Carnivore | *Ca* | For species eating other animals. | 481 |
| Crawls | *Cr* | Whether the species crawls. | 184 |
| Cyanobacteria | *Cy* | Member of the cyanobacteria phylum. | 1 |
| Detritivore | *De* | For species eating detritus. | 355 |
| Detritus | *Ds* | Whether the species can be classifying as a detritus. | 2 |
| Fungivore | *Fg* | For species eating fungi. | 111 |
| Fungi | *Fu* | Member of the fungi kingdom. | 2 |
| HasShell | *HS* | Whether the species has a shell. | 274 |
| Herbivore | *He* | For species eating plants. | 130 |
| Immobile | *Im* | For immobile species. | 85 |
| IsHard | *IH* | Whether the species has a tough exterior (but not a shell). | 418 |
| Jumps | *Ju* | Whether the species can jump. | 30 |
| LongLegs | *LL* | For species with long legs. | 59 |
| Mollusca | *Mo* | Member of the mollusca phylum. | 45 |
| Nematoda | *Ne* | Member of the nematoda phylum. | 5 |
| Plantae | *Pl* | Member of the plant kingdom. | 3 |
| Protozoa | *Pr* | Member of the protozoa kingdom. | 3 |
| ShortLegs | *SL* | For species with short legs. | 538 |
| UsePoison | *UP* | Whether the species uses poison. | 177 |
| WebBuilder | *WB* | Whether the species builds webs. | 89 |
| Body mass | *M* | Natural logarithm of the body mass in grams | 881 |
| $Ph_0$ | $Ph_0$ | Coordinate on the first axis of a PCA of phylogenetic distances | 881 |
| $Ph_1$ | $Ph_1$ | Coordinate on the second axis of a PCA of phylogenetic distances | 881 |

### 2.2 *K*-nearest neighbour

Both our recommendation and matrix imputation approaches use the *K*-nearest neighbour (*K*NN) algorithm [24]. The *K*NN algorithm is an instance-based method, it does not build a general internal model of the data, but instead tries to fill missing entries by a majority vote based on the *K* nearest (i.e. most similar) entries given some distance metrics. In the case of recommendation, there is no concept of “missing entry”, each species is described by a set of traits and a set of preys, and the algorithm will recommend new preys to the species based on the preys of its *K* nearest neighbours. For example, if *K* = 3, we take the set of preys of the three most similar species to decide which prey to recommend. If species *A* is found twice and *B* once in the set of preys of the most similar species, we will recommend *A* first (assuming, of course, that the species does not already have this prey). See table 3 for a complete example of recommendation. In the “Netflix” problem, this is equivalent to recommend new TV series/movies to a user by searching for the users with the most similar taste and using what they liked as recommendation. It is also possible to tackle the reverse problem: Amazon uses item-based recommendations, in which case we are looking for similar items instead of similar users to base our recommendations [2].

**Table 3:** Fictional example to illustrate recommendations with *K* nearest neighbour using the Tanimoto distance measure modified to include species traits. We are trying to recommend a prey to species 0 given that the three most similar species are species 6, 28, and 70. For example, the distance to species 70 would be *w*_t_0.5 + (1 - w_t_)1/3. To find recommendations, the set of preys found in the *K* = 3 most similar entries is computed, in this case {812 = 2, 70 = 2, 72 = 1}, leading to the list of recommendations [812, 70, 72]. Because they are found most often in the K most similar species, candidates 812 and 70 will be suggested before 72. To test this approach, we remove a prey from a species and check whether the algorithm recommend the missing prey. Especially with low *K*, it’s possible that no recommendations can be found, for example if the most similar species has the exact same preys.

| Species ID | Traits | Preys | Most similar | Recommendations |
| --- | --- | --- | --- | --- |
| 0 | $\{Ar, Ca\}$ | $\{6, 42, 47\}$ | $\{6, 28, 70\}$ | $[812, 70, 72]$ |
| 6 | $\{Ar, Ca\}$ | $\{42, 47, 70, 72\}$ | | |
| 28 | $\{Ar, Ca\}$ | $\{42, 47, 70, 812\}$ | | |
| 70 | $\{Ca\}$ | $\{42, 47, 812\}$ | | |
| ... | ... | ... |  |  |

For matrix imputation, if we want to know if a species preys on *A*, we look at how many of the *K* most similar species prey on *A* (a ratio that can be interpreted as a probability). In this case, the problem is seen as a matrix with missing entries, so the question is not to recommend new preys to a predator, but whether a specific relationship exists (see figure 1 for a complete example). It is often suggested to avoid picking *K* that are multiple of the number of classes to avoid ties [29]. Here there are two classes: interaction and non-interactions, so we will only use odd *K*s.

**Figure 1:**
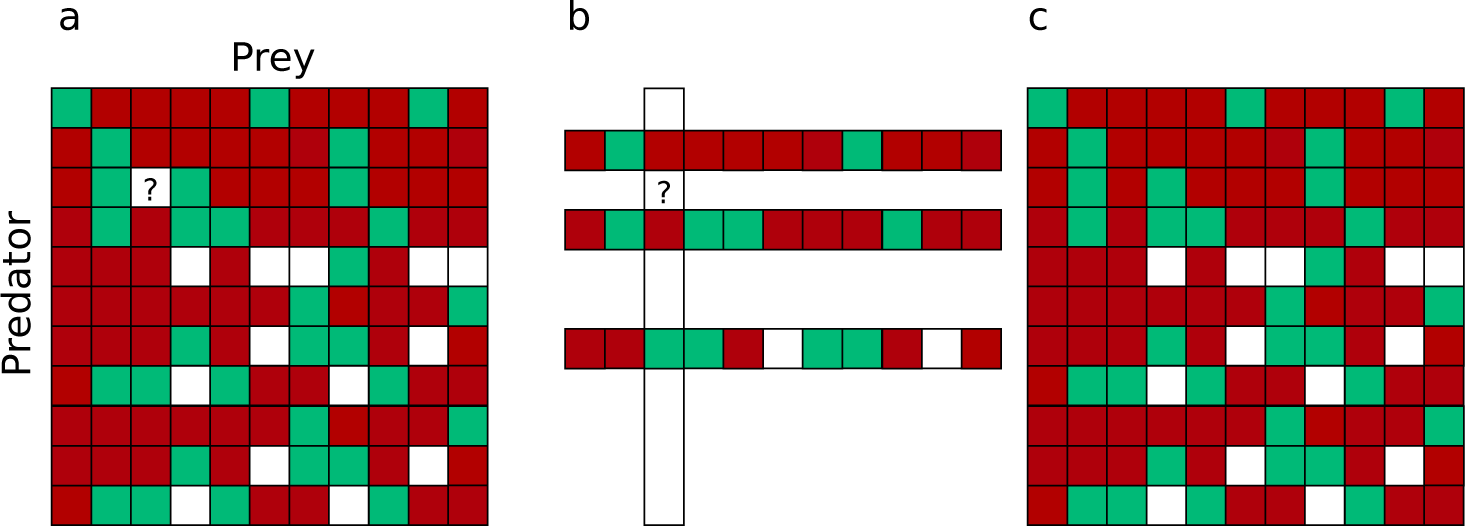
**A**: The initial matrix with missing entries, green squares are used for interactions, red for non-interactions, and white for missing values. Rows represent entries and columns features. In our case, we have a square matrix where rows are species and the columns represent their preys. We want to find the value denoted by **X**. **B**: The K-nearest neighbours algorithms look at the entries that are the most similar to the entry with the missing value, pick the 3 closest (*K*= 3), and use the values at the columns with the missing entry. **C**: In this case, for the column with the missing entry, the 3 nearest neighbours have 2 non-interactions and 1 interaction, so the algorithm fills the entry with a non-interaction.

Different distance measures can be used. Here, we will use the Tanimoto coefficient for recommendations and the Euclidean distance for matrix imputation. Choosing the right value for *K* is tricky. Low values give high importance to the most similar entries, while high values provide a larger set of examples. Fortunately, the most computationally intensive task is to compute the distances between all pairs, a step that is independent of *K*. As a consequence, once the distances are computed, we can quickly run the algorithm with different values of *K*.

### 2.3 Recommendation

The Tanimoto (or Jaccard) similarity measure is defined as the size of the intersection of two sets divided by their union, or:

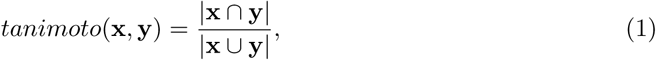

Since it is a similarity measure in the [0,1] range, we can transform it into a distance function with 1 – *tanimoto*(x,y). The distance function will use two types of information: the set of traits of the species (see table 2) and their set of preys. We define the distance function with traits as:

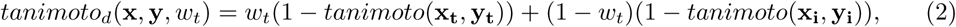

where *w_t_* is the weight given to traits, x_t_ and y_t_ are the sets of traits for species *x* and *y*, and x_i_, y_i_ are their sets of preys. Thus, when *w_t_* = 0, only interactions are used to compute the distance, and when *w_t_* = 1, only traits are used. See table 3 for an example.

The data is the set of preys and binary traits for each species (Table 2). To test the approach, we randomly remove an interaction for each species and ask the algorithm to recommend up to 10 preys for the species with the missing interaction. We count how many recommendations are required to retrieve the missing interactions and compute the top1, top5, and top10 success rates, which are defined as the probabilities to retrieve the missing interaction with 1, 5, or 10 recommendations. We repeat this process 10 times for each species with at least 2 preys (7200 attempts). We test all odd values of *K* from 1 to 19, and *w_t_* = {0,0.2, 0.4,0.6, 0.8,1}. We also divided species in groups according to the number of preys they have to see if it is easier to find the missing interaction for species with fewer preys.

### 2.4 Matrix imputation

The *K*NN algorithm with Euclidean distance works with both positive and negative entries. In this case, an interaction is represented with a value of 1, while a non-interaction is a represented with 0, in a *n* × *n* matrix (*n* = 881). The goal is to predict the value of a missing entry (Figure 1). The Euclidean distance is defined as

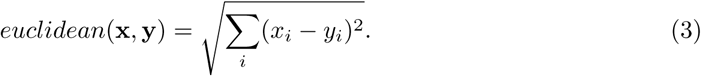

However, we want to give different weights to different aspects of the species, so we compute the distance between two species as:

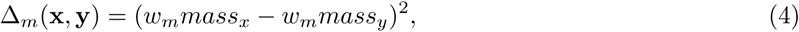

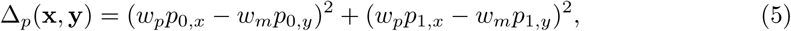

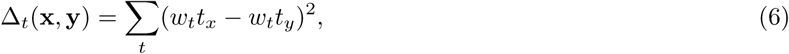

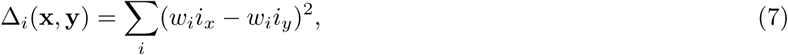

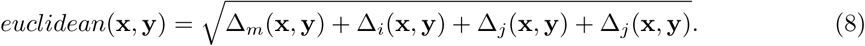

Where *w_m_*, *w_p_, w_t_, w_i_* are the weight given to body mass, the two coordinates of a classical multidimensional scaling, binary traits, and interactions, respectively. For simplicity we require that *w_m_* + *w_p_* + *w_t_* + *w_i_* = 1.

The data is an 881 × 881 interaction matrix. To test the *K*NN algorithm with the Euclidean distance, we randomly remove a single interaction from the matrix, ask the algorithm to fill the entry, and count how many times the correct value is retrieved. For each set of parameters tested, we repeat this process 50 000 times, and count the number of true positives (tp), true negatives (tn), false positives (fp) and false negatives (fn). The score for predicting interactions (*Score_y_*), non-interactions (*Score_-y_*) and the accuracy are defined as

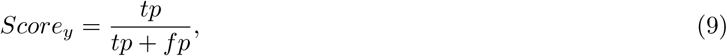

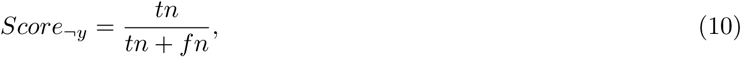

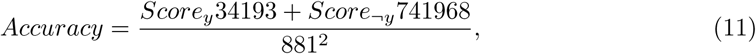

with 34193 and 741968 being the number of observed interactions and non-interactions in the 881 by 881 matrix. We also use the True Skill Statistics (TSS), defined as

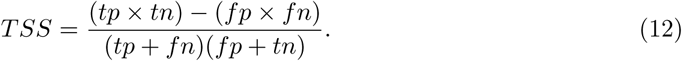

The *TSS* ranges from −1 to 1.

### 2.5 Supervised learning

We also do a simple test with random forests to see if it is possible to predict interactions in this data-set using only the traits [6]. In this case, the random forests perform supervised learning: we are trying to predict *y* (interaction) from the vector of traits **x** by first learning a model on the training set, and testing the learned model on a testing set. We keep 5% of the data for testing. We perform grid search to find the optimal parameters for the random forests.

### 2.6 Code and Data

Since several machine learning algorithms depends on computing distances (or similarities) for all pairs, many data structures have been designed to compute them efficiently from kd-trees discovered more than thirty years ago [12] to ball trees, metric skip lists, navigating nets [20], and cover trees [5, 20]. We use an exact but naive approach that works well with small datasets. Since *distance*(*x,y*) = 0 if *x* = *y* and *distance*(*x,y*) = *distance*(*y,x*), our C++ implementation stores the distances in a lower triangular matrix without the diagonal, yielding *n*(*n* − 1)/2 distances to compute. We used Scikit for random forests [25]. The C++11 code for the *K*NN algorithm, Python scripts for random forests, and all data-sets used are available at https://github.com/PhDP/EcoInter [10].

## 3 Results

### 3.1 Recommendation

While matrix imputation has a 50% change of success by random, the Tanimoto *K*NN needs to pick the right prey among up to 881 possibilities. Yet, it succeeds on its first recommendation around 50% of the times. When the first recommendation fails, the next 9 recommendations only retrieve the right species around 15% of the times so the top5 and top10 success rates are fairly close to the top1 success rate (see figure 2). The Tanimoto measure is particularly effective for species with fewer preys, achieving more than 80% success rate for species with 10 or fewer preys (Figure 3).

**Figure 2:**
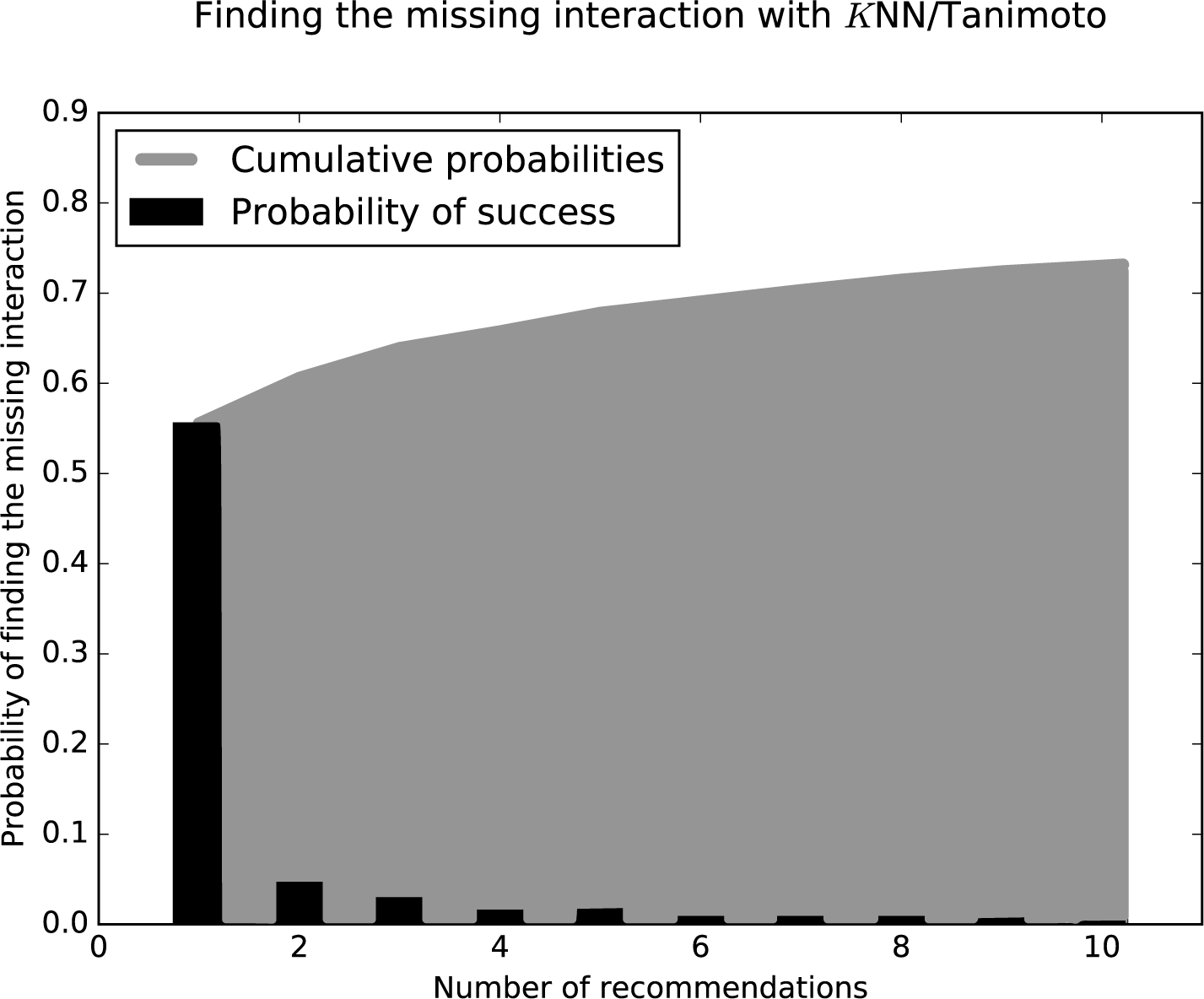
After removing a prey from a predator, we ask the KNN algorithms with Tanimoto measure to make 10 recommendations (from best to worst). The figure shows how many recommendations are required to retrieve the missing interaction. Most retrieved interactions are found with the first attempt. This data was generated with *K* = 7 and *w_t_* = 0.

**Figure 3:**
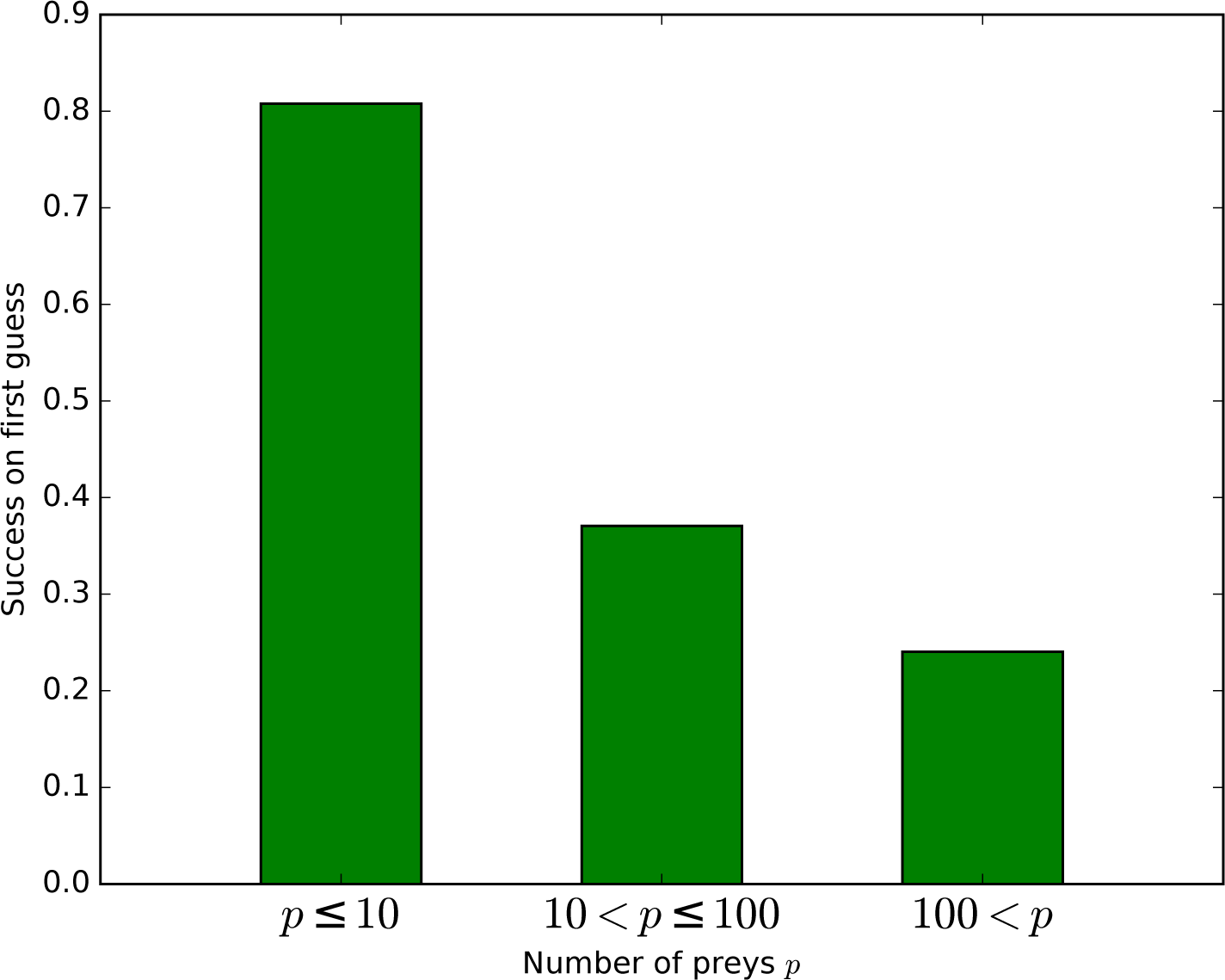
Success on first guess with Tanimoto similarity as a function of the number of prey. The KNN algorithm with Tanimoto similarity is more effective at predicting missing preys when the number of preys is small. This is probably in good part because there are more information available to the algorithm, since 473 species have 10 or fewer preys, 295 have between 10 and 100, 103 species have more than 100 preys.

The highest first-try success rates (the probability to pick the missing interaction on the first recommendation) are found with *K* = 7 and no weights to traits, and with *K* = 17 and a small weight of 0.2 to traits (Table 4). Overall, the value of *K* had little effect on predictive ability.

**Table 4:**
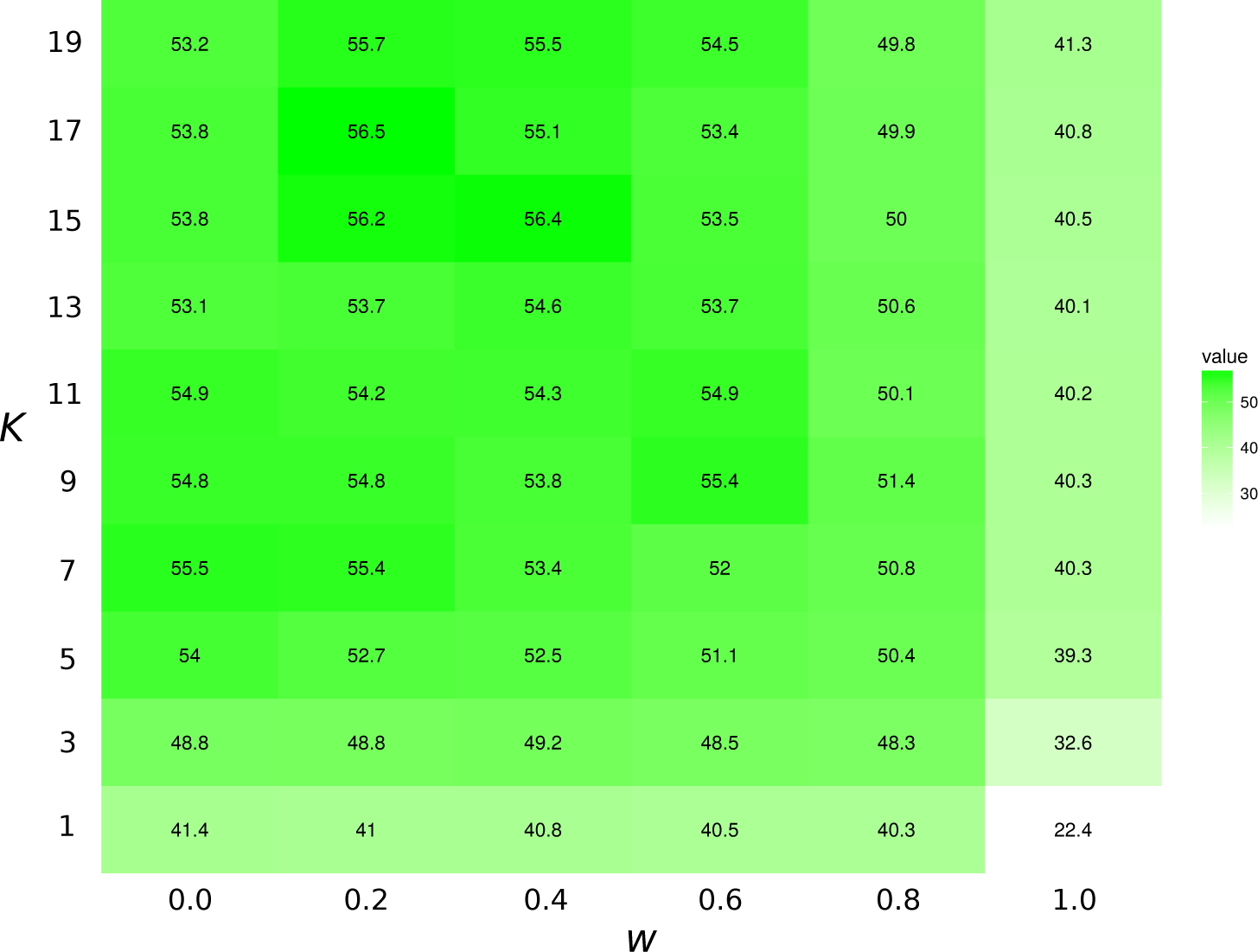
Top1 success rates for the *K*NN/Tanimoto algorithm with various *K* and weights to traits. When *w_t_* = 0.0, the algorithm will only use interactions to compute similarity between species. When *w_t_* = 1, the algorithm will only consider the species’ traits (see table 2). The value is the probability to retrieve the correct missing interaction with the first recommendation. For each entry, *n* = 871 (the number of species minus 10, the number of species with no preys). The best result is achieved with *K* = 17 and *w* = 0.2, although the results for most values of *K* and w = [0.0, 0.2] are all fairly close. The success rate increases with *K* when only traits are considered (*w* = 1).

### 3.2 Matrix imputation

We show our results with *K* Nearest neighbours algorithm in table 5. The best result is achieved with *K* = 1 but has a TSS of only 0.66 (Table 5). More than 99% of non-interactions are predicted correctly, but only 2/3 of the interactions are predicted correctly (Table 5).

**Table 5:** Matrix imputation with Euclidean distance. This tables uses the weights *w_m_* = 1 /3, *w_p_* = 0, *w_t_* = 1 /3, *w_i_* = 1/3.

| K | $Score_y$ | $Score_{\neg y}$ | Accuracy | TSS |
| --- | --- | --- | --- | --- |
| 1 | 0.6726 | 0.9939 | 0.9804 | 0.6664 |
| 3 | 0.6277 | 0.9962 | 0.9807 | 0.6239 |
| 5 | 0.5671 | 0.9975 | 0.9794 | 0.5645 |
| 7 | 0.5232 | 0.9978 | 0.9780 | 0.5210 |
| 9 | 0.4754 | 0.9984 | 0.9765 | 0.4739 |

As for the weights *w_m_* (body mass), *w_p_* (coordinates from a classical multidimensional scaling of the phylogeny), *w_t_* (binary traits), *w_i_* (interactions), the optimal values for *w_m_* and *w_i_* lie between 1/3 and 1/4 and vary a bit with different values of *K*. *w_t_* has a small but consistent effect: changing the weight from 0 to 1/3 improves the result by roughly 1%, but higher values will start to decrease predictive ability. Positive values of *w_p_* always have a negative effect on predictive abilities. With minor variations with *K*, the optimal weights are thus *w_m_* ≈ 1/3, *w_p_* = 0, *w_t_* = 1/3, *w_i_* ≈ 1/3. The optimal weight to *w_i_* increases with *K*.

### 3.3 Supervised learning

Random forests predict correctly 99.55% of the non-interactions and 96.81% of the interactions, for a TSS of 0.96. Much of this accuracy is due to the three real-valued traits (body mass, *Ph*_0_, *Ph*_1_). Without them, too many entries have the same feature vector **x**, making it impossible for the algorithm to classify them correctly. Removing the binary traits has little effect on the model. With only body mass, *Ph*_0_, *Ph*_1_, the TSS of the random forests is 0.94.

## 4 Discussion

We applied different machine learning techniques to the problem of predicting binary species interactions. Recommendation is arguably a better fit for binary species interactions, since it is essentially the same problem commercial recommenders such as Netflix face: given that a user like item *i*, what is the best way to select other items the user would like? In this case, users are species, and the items are their preys, but the problem is the same. In both cases, we can have solid positive evidence (observed or implied interactions), but rarely have proofs of non-interactions. The approach yields strong results, with a top1 success rate above 50% in a food web with up to 881 possibilities. The approach could be used, for example, to reconstruct entire food webs using global database of interactions [27]. The method’s effectiveness rely on nestedness: how much species cluster around the same set of preys in a food web [16]. Thus, it should be less effective in food webs with more unique predators.

The *K*NN algorithm falls into the realm of unsupervised learning, where the goal is to find patterns in data [24]. The other class of machine learning algorithms, supervised learning, have the clearer goal of predicting a value y from a vector of features **x**. For example, in supervised learning, we would try to predict an interaction y from the vector of traits **x**, while our unsupervised approach allow us to fill entries from an incomplete matrix regardless of what the entry is (interaction or trait). For matrix imputation, the *K*NN yields less than impressive results, but our random forests test on the same data-set achieves a TSS of 0.96 using the binary traits, body mass, and the coordinates of the multidimensional scaling. A random forest can build effective predictive models by creating complex rules based on the traits, while the *K*NN algorithm relies on a simplistic distance metrics. However, the *K*NN approach has some advantages over supervised learning, namely the capacity to fill any entries from an interaction matrix and use the information from other species’ interactions. The solution is likely to *learn* distance metrics [4] instead of using a fixed formula. This would allow complex rules while maintaining the *K*NN’s ability to fill arbitrary matrices.

Learning distance metrics is a promising avenue to improve our results. Much efforts on the Netflix prize focused on improving similarity measures [30, 19], and custom similarity metrics can be used to improve unsupervised classification algorithms [4]. Learning distance metrics from data is a common way to improve methods based on a nearest neighbour search [34, 4], allowing the measure itself to be optimized. We only used the *K* nearest neighbour algorithm for unsupervised learning, but several other algorithms can be used to solve the “Netflix problem”. For example: techniques based on linear programming, such as recent exact methods for matrix completion based on convex optimization [8] or low-rank matrix factorization. The latter method reduces a matrix to a multiplication between two smaller matrices, which can be used both to predict missing entries and to compress large matrices into small, more manageable matrices [31]. Given enough data, deep learning methods such as deep Boltzmann machines could also be used [35]. Deep learning revolutionized machine learning with neural networks made of layers capable of learning increasingly detailed representations of complex data [18]. Many of the most spectacular successes of machine learning use deep learning [21]. However, learning several neural layers to form a deep networks would require larger data-sets.

The low sensibility to *K* in recommendations compared to imputation is interesting. This is cause by the fact that, as *K* grows, the set of species includes more and more unrelated species with widely different set of preys. For imputation, adding more species with different preys means it is likely to misclassify an interactions as a non-interaction. However, if we increase *K* from k to *k* + *δ* for a recommendation, the species in *δ* range are not only less similar, but they are less likely to share preys among themselves. Since recommendations are based on how many times a prey is found in the *K* nearest species, the species in the *δ* range are unlikely to have as much weight as the first *k* species.

Our results have two limitations. It is possible that our food web was exceptionally simple, and that a food web with distinct structural properties would behave differently, especially if it has lower nestedness. The success of the *K*NN algorithms depends on local structure: how much can we learn from similar species. If each species has a unique set of preys, the *K*NN will struggle more. Also, a deeper issue is that real food webs are not binary structures. Species, populations, and individuals have different densities, prey more strongly on some resources than others, and have preferences. In a binary matrix, we can predict if two species will interaction while completely ignoring the rest of the network, but real food webs involve complex indirect relationships [33]. It is unclear how much we can learn about ecosystems and species interactions from binary matrices, and our results show that binary interactions are mostly independent of the community, since we are able to effectively predict if two species interactions given only three traits. Species interactions are better represented with a weighted hypergraph [13], which are well-suited to model relations with an arbitrary number of participants, where the hyperedge would allow for complex indirect relationships to be included. Understanding these hypergraphs is outside the scope of the *K*NN algorithm but could be understood with modern techniques such as Markov logic [28].

Recommendation (*K*NN algorithm with Tanimoto distance) and supervised learning (random forests) are complementary techniques. Supervised learning is more useful when we have traits and no information about interactions, but it is useless without the traits. On the other hand, the recommender performs well without traits but requires at least partial information about interactions, although it might be possible to use the interactions from different food webs. Matrix imputation might provide the best of the both worlds, allowing us to use both traits and species interactions, but the distance metrics we used performed poorly and we suggest more research could be done on developing better distance metrics for ecological interactions, or learning these metrics from data.

## 5 Acknowledgements

PDP has been funded by an Alexander Graham Bell Graduate Scholarship from the National Sciences and Engineering Research Council of Canada, an Azure for Research award from Microsoft, and benefited from the Hardware Donation Program from NVIDIA. We tank the team of U. Brose for documenting the food webs. DG is funded by the Canada Research Chair program and NSERC Discovery grant. TP is funded by an NSERC Discovery grant and an FQRNT Nouveau Chercheur grants.

## References

[1] A Aderhold, D Husmeier, JL Lennon, CM Beale, and VA Smith. Hierarchical bayesian models in ecology: Reconstructing species interaction networks from non-homogeneous species abundance data. Ecological Informatics, 11:55–64, 2012.

[2] CC Aggarwal. Recommender Systems. Springer, 2016.

[3] I Bartomeus, D Gravel, J Tylianakis, M Aizen, I Dickie, and M Bernard-Verdier. A common framework for identifying linkage rules across different types of interactions. Functional Ecology, 2016.

[4] A Bellet, A Habrard, and M Sebban. Metric Learning. Morgan & Claypool, 2015.

[5] A Beygelzimer, S Kakade, and J Langford. Cover trees for nearest neighbor. In Proceedings of the 23nd International Conference on Machine Learning, 2006.

[6] L Breiman. Random forests. Machine Learning, 45(1):5–32, 2001.

[7] EF Canard, N Mouquet, D Mouillot, M Stanko, D Miklisova, and D Gravel. Empirical evaluation of neutral interactions in host-parasite networks. American Naturalist9, 183:468–479, 2014.

[8] EJ Candès and B Recht. Exact matrix completion via convex optimization. Foundations of Computational mathematics, 9(6):717–772, 2009.

[9] TF Cox and MAA Cox. Multidimensional Scaling. Chapman and Hall, 2001.

[10] P Desjardins-Proulx. github.com/phdp/articles. http://doi.org/10.5281/zenodo.161602. 2016.

[11] C Digel, A Curtsdotter, J Riede, B Klarner, and U Brose. Unravelling the complex structure of forest soil food webs: higher omnivory and more trophic levels. In Oikos, volume 123, pages 1157–1172, 2014.

[12] JH Friedman, JL Bentley, and RA Finkel. An algorithm for finding best matches in logarithmic expected time. Transactions on Mathematical Software, 3(3):209–226, 1977.

[13] J Gao, Q Zhao, W Ren, A Swami, R Ramanathan, and A Bar-Noy. Dynamic shortest path algorithms for hypergraphs. Modeling and Optimization in Mobile, Ad Hoc and Wireless Networks, pages 238–245, 2012.

[14] AJ Golubski, EE Westlund, J Vandermeer, and M Pascual. Ecological networks over the edge: Hypergraph trait-mediated indirect interaction (tmii) structure. Trends in Ecology and Evolution, 31(5):1083–1090, 2016.

[15] D Gravel, T Poisot, C Albouy, L Velez, and D Mouillot. Inferring food web structure from predator-prey body size relationships. Methods in Ecology and Evolution, 4(11):1083–1090, 2013.

[16] PR Guimaraes and P Guimaraes. Improving the analyses of nestedness for large sets of matrices. Environmental Modelling and Software, 21(10):1512–1513, 2006.

[17] A Halevy, P Norvig, and F Pereira. The unreasonable effectiveness of data. IEEE Intelligent Systems, 24:8–12, 2009.

[18] GE Hinton, S Osindero, and YW Teh. A fast learning algorithm for deep belief nets. Neural computation, 18(7):1527–1554, 2006.

[19] T Hong and D Tsamis. Use of KNN for the Netflix Prize. 2006.

[20] M Izbicki and CR Shelton. Faster cover trees. In Proceedings of the 32nd International Conference on Machine Learning, 2015.

[21] V Mnih, K Kavukcuoglu, D Silver, A Graves, I Antonoglou, D Wierstra, and M Riedmiller. Playing atari with deep reinforcement learning. arXiv, 2013.

[22] I Morales-Castilla, MG Matias, D Gravel, and MB. Araújoemail. Inferring biotic interactions from proxies. Ecological Informatics, 30(6):347–356, 2015.

[23] N Mouquet, V Devictor, CN Meynard, F Munoz, LF Bersier, J Chave, P Couteron, A Dalecky, C Fontaine, and D Gravel. Ecophylogenetics: advances and perspectives. Biological reviews, 87(4):769–785, 2012.

[24] KP Murphy. Machine Learning: A Probabilistic Perspective. The MIT Press, 2012.

[25] F. Pedregosa, G. Varoquaux, A. Gramfort, V. Michel, B. Thirion, O. Grisel, M. Blondel, P. Prettenhofer, R. Weiss, V. Dubourg, J. Vanderplas, A. Passos, D. Cournapeau, M. Brucher, M. Perrot, and E. Duchesnay. Scikit-learn: Machine learning in Python. Journal of Machine Learning Research, 12:2825–2830, 2011.

[26] SL Pimm. Food Webs. Springer, 1982.

[27] JH Poelen, JD Simons, and CJ Mungall. Global biotic interactions: An open infrastructure to share and analyze species-interaction datasets. Ecological Informatics, 24:148–159, 2014.

[28] M Richardson and P Domingos. Markov logic networks. Machine Learning, 62(1-2):107–136, 2006.

[29] S Theodoridis. Machine Learning: A Bayesian and Optimization Perspective. Academic Press, 2015.

[30] A Toscher and M Jahrer. The BigChaos solution to the Netflix prize. 2008.

[31] RJ Vanderbei. Linear programming: Foundations and extensions. 2013.

[32] RJ Williams and ND Martinez. Simple rules yield complex food webs. Nature, 404:180–183, 2000.

[33] JT Wootton. The nature and consequences of indirect effects in ecological communities. Annual Review of Ecology and Systematics, pages 443–466, 1994.

[34] EP Xing, AY Ng, MI Jordan, and S Russell. Distance metric learning with application to clustering with side-information. Advances in neural information processing systems, 15:505–512, 2003.

[35] J Zhang. Deep transfer learning via restricted boltzmann machine for document classification. In ICMLA: Machine Learning and Applications, volume 1, pages 323–326, 2011.

